# “*Candidatus* Siderophilus nitratireducens”: a psychrophilic, *nap*-dependent nitrate-reducing iron oxidizer within the new order Siderophiliales

**DOI:** 10.1101/2023.09.04.556225

**Authors:** Francesc Corbera-Rubio, Gerben R. Stouten, Jantinus Bruins, Simon F. Dost, Alexander Y. Merkel, Simon Müller, Mark C. M. van Loosdrecht, Doris van Halem, Michele Laureni

## Abstract

Nitrate leaching from agricultural soils is increasingly found in groundwater, a primary source of drinking water worldwide. This nitrate influx can potentially stimulate the biological oxidation of iron in anoxic groundwater reservoirs. Nitrate-reducing iron-oxidizing (NRFO) bacteria have been extensively studied in laboratory settings, yet their ecophysiology in natural environments remains largely unknown. To this end, we established a pilot-scale filter on nitrate-rich groundwater to elucidate the structure and metabolism of nitrate-reducing iron-oxidizing microbiomes under oligotrophic conditions mimicking natural groundwaters. The enriched community stoichiometrically removed iron and nitrate consistently with NRFO metabolism. Genome-resolved metagenomics revealed the underlying metabolic network between the dominant iron-dependent denitrifying autotrophs and the less abundant organoheterotrophs. The most abundant genome belonged to a new *Candidate* order, named Siderophiliales. This new species, “*Candidatus* Siderophilus nitratireducens”, carries central genes to iron oxidation (cytochrome *c cyc2*), carbon fixation (*rbc*), and for the sole periplasmic nitrate reductase (*nap*). To our knowledge, this is the first report of *nap*-based lithoautotrophic growth, and we demonstrate that iron oxidation coupled to dissimilatory reduction of nitrate to nitrite is thermodynamically favourable under realistic Fe^3+^/Fe^2+^ and 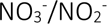 concentration ratios. Ultimately, by bridging the gap between laboratory investigations and real-world conditions, this study provides insights into the intricate interplay between nitrate and iron in groundwater ecosystems, and expands our understanding of NRFOs taxonomic diversity and ecological role.

## 1. Introduction

Globally, approximately one third of the nitrogen applied to agricultural soils is lost via leaching to the surrounding waterbodies (1). This has led to elevated nitrate (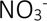) levels in anoxic groundwaters, a primary source of drinking water worldwide (2). Owing to population growth and agriculture intensification, nitrate concentrations in subsurface waters are expected to continue increasing (3). Besides its direct impact on human health (4), nitrate can significantly alter the biogeochemistry of groundwater reservoirs (5). Nitrate promotes the oxidation of sulfide and in particular of iron (Fe) – the most prevalent groundwater contaminant – leading to the mobilisation of heavy metals and the emission of greenhouse gases (6). Despite these implications, the consequences of nitrate-iron interactions on ecosystems and drinking water production systems remain largely unexplored. A detailed understanding of the underlying principles is paramount for anticipating and mitigating current and future challenges, as well as for exploring potential synergies and biotechnological opportunities.

Nitrate-reducing iron-oxidizing (NRFO) bacteria couple the anoxic reduction of nitrate to the oxidation of Fe^2+^ (eq.1). Since their discovery in 1996 by Straub et al. (1996), NRFO microorganisms have been the focus of extensive research both in pure and mixed cultures (reviewed in (8)), and several complete genomes are already publicly available (9,10). The metabolic versatility of NRFO bacteria spans from lithoautotrophic to mixotrophic growth (11), to partial denitrification using nitric oxide (NO) (12) and nitrous oxide (N_2_O) (9) as terminal electron acceptors. At the same time, due to the inherently low energetic yield of iron oxidation, NRFO bacteria live close to the thermodynamic edge (13). Their fitness is highly dependent on environmental factors such as substrate and product availability, pH and temperature (14), and chemical reactions - such as the quasi-instantaneous precipitation of the biologically formed Fe^3+^ - can play a pivotal role by modulating iron and nitrogen concentrations (15). However, our current understanding is largely based on laboratory settings, and does not necessarily reflect the complexity of natural and engineered ecosystems where several (a)biotic reactions occur simultaneously at temperatures significantly lower than tested to date (16).

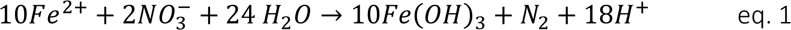

To address these knowledge gaps, we established a pilot-scale filter on anoxic groundwater containing both Fe^2+^ and 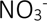. The emulated groundwater conditions allowed for the establishment of a microbial enrichment that simultaneously removed Fe^2+^ and 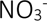. In depth metagenomic analysis of the steady-state community revealed a new order-level NRFO lineage, deepening our understanding of their taxonomic diversity and ecological roles. Overall, our study bridges the gap between laboratory studies and real-world conditions, and offers a nuanced view on the intricate interplay between nitrate and iron in groundwater ecosystems.

## 2. Results

### 2.1. Nitrate-dependent iron removal irrespective of the limiting nutrient

Stable anoxic nitrate and iron removal was achieved after less than three weeks of operation and maintained for over 100 days in a pilot-scale filter fed with nitrate-rich anoxic groundwater (Figure S3). With nitrate as the limiting nutrient at both groundwater (8.9 ± 2.8 μM) and nitrate-amended concentrations (13.5 ± 1.5 and 20.2 ± 2.4 μM), effluent nitrate concentrations were consistently below detection limit (1 μM). Throughout the nitrate-limiting period, 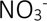 and Fe^2+^ were consumed at a 7.1 ± 1.4 Fe^2+^: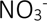 molar ratio (Figure 1Figure 1). Oxygen was always below the quantification limit of 3 μM, and roughly 80 μC-mol dissolved organic carbon (DOC·l^−1^) was consistently removed from the influent, likely due to the formation of Fe^2+^-DOC complexes formation owing to the non-biodegradable nature of organic matter in Dutch groundwater (17). Ammonia consumption was negligible (<0.1 µM). Effluent nitrite concentrations stayed below the detection limit (<0.2 μM), while other denitrification intermediates – nitric oxide and nitrous oxide – were not measured. The observed consistent stoichiometric coupling between nitrate and iron removals strongly suggests Fe^2+^ oxidation to be primarily driven by microbial nitrate-reducing iron oxidation.

**Figure 1.**
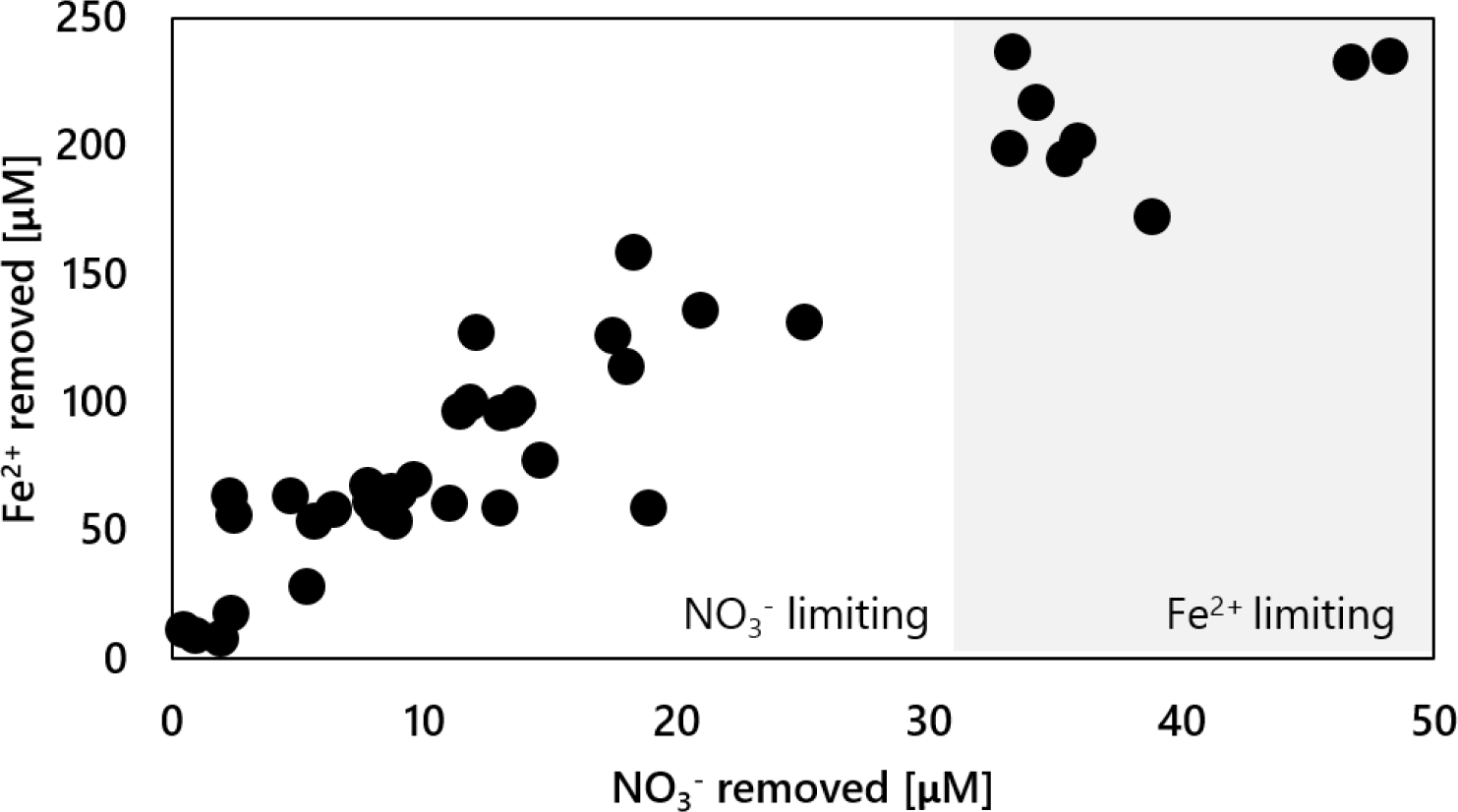
Simultaneous 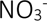 and Fe^2+^ removals in the groundwater-fed pilot-scale filter during the 120 days of continuous operation. The groundwater Fe^2+^ concentration was constant throughout the experiment (236 ± 4 µM). 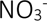 was dosed in the influent to step-wise increase the natural groundwater concentration from 8.1 ± 2.1 to 20.2 ± 2.4 µM (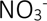 limitation), and up to 83.8 ± 0.6 µM (Fe^2+^ limitation).

### 2.2. Microbial community dominated by iron oxidizers and denitrifiers

Metagenomic DNA sequencing yielded a total of 107,512 and 8,754,261 quality filtered short and long reads, respectively. After assembly and polishing, this resulted in 19,127 contigs with an N50 value of 15,927. Contigs binning resulted in 13 high and medium quality metagenome assembled genomes (MAGs) as defined by (18) with a relative abundance exceeding 0.5% of the quality filtered long reads. Collectively, these 13 most abundant genomes accounted for 66.9% of the total quality filtered reads, and belonged to four phyla: *Proteobacteria* (51.6%), *Actinobacteria* (8.3%), *Bacteroidetes* (5.6%) and *Chloroflexi* (1.4%) (Table 1). Every genome in the community contained at least one denitrification enzyme genes, while five featured the genetic potential for iron oxidation. Notably, all putative iron oxidizers also possessed the genetic repertoire for carbon fixation. The most abundant MAG (MAG.13), accounting for 19.3% of the community, could only be taxonomically classified at class level (*Gammaproteobacteria*). Given its high abundance and potential metabolic relevance, the taxonomy and metabolic potential of MAG.13 was further investigated (Table S2).

**Table 1.**
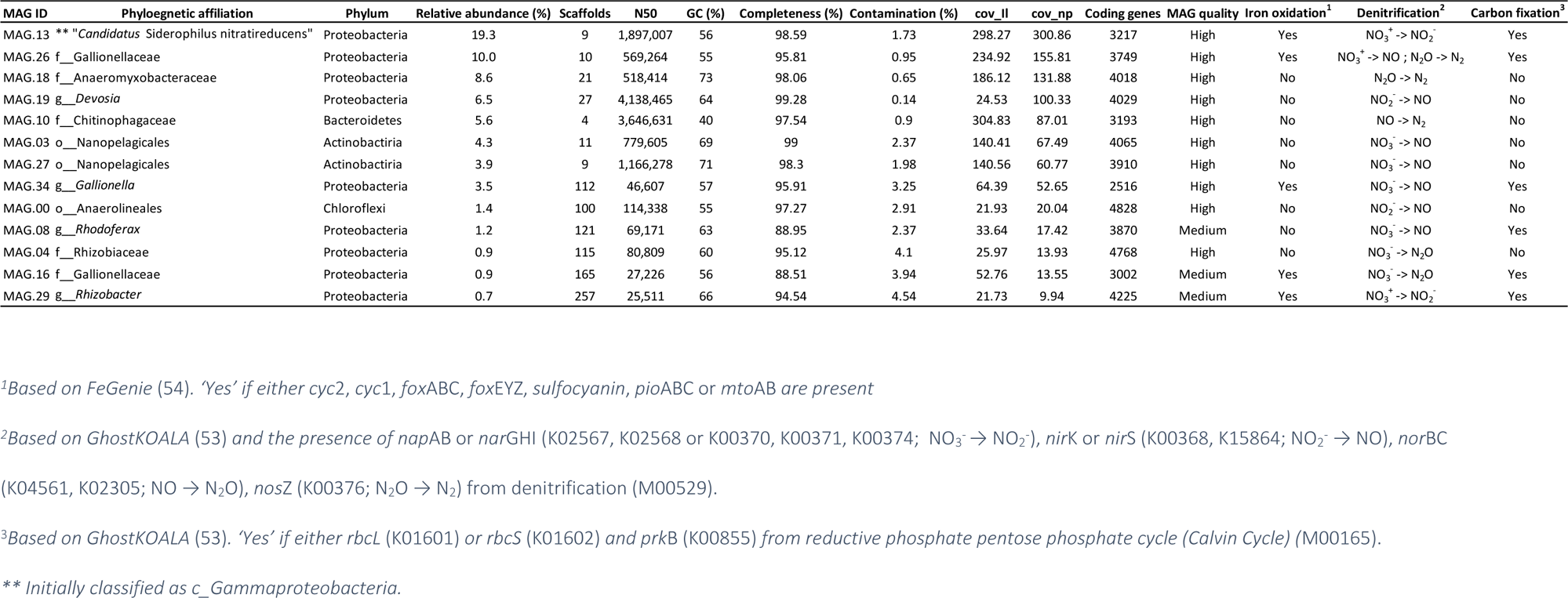
General characteristics of the MAGs recovered from the pilot-scale filter. The last three columns indicate the presence/absence of the essential genes for iron oxidation, denitrification and carbon fixation via the reductive phosphate pentose phosphate cycle. Cov_il and cov_np are the coverage with Illumina short reads and Nanopore long reads, respectively.

### 2.3. “Candidatus *Siderophilus* nitratireducens” represents a new order within Gammaproteobacteria

Our phylogenomic analysis based on the concatenated amino acid sequences of 120 bacterial single copy conservative marker genes revealed that MAG.13 belongs to a bacterium forming a new order-level lineage Ga0077554 (GTDB release 08-RS214) within the class *Gammaproteobacteria,* with no known closely related pure-culture representatives (Figure 2). We propose to name the new species “*Candidatus* Siderophilus nitratireducens” gen.nov., sp.nov., a member of the *Candidate* order and family Siderophiliales and Siderophiliaceae, respectively. This lineage, along with several other MAGs from similar groundwater habitats (19), is mostly related to lineages including lithoautotrophic sulfur-ozidizing bacteria from the genera *Sulfuriflexus*, *Thioalbus* and the members of the order *Thiohalomonadales*, including *Thiohalomonas*, *Sulfurivermis* and *Thiohalophilus*. Average nucleotide identity and DNA-DNA hybridization comparison between *Ca.* Siderophilus nitratireducens and its closest relative, a MAG from a drinking water treatment plant (GCA_001464965.1), indicated that the two organisms belong to the same genus but different species (ANI = 89%, DDH = 36.6%).

**Figure 2.**
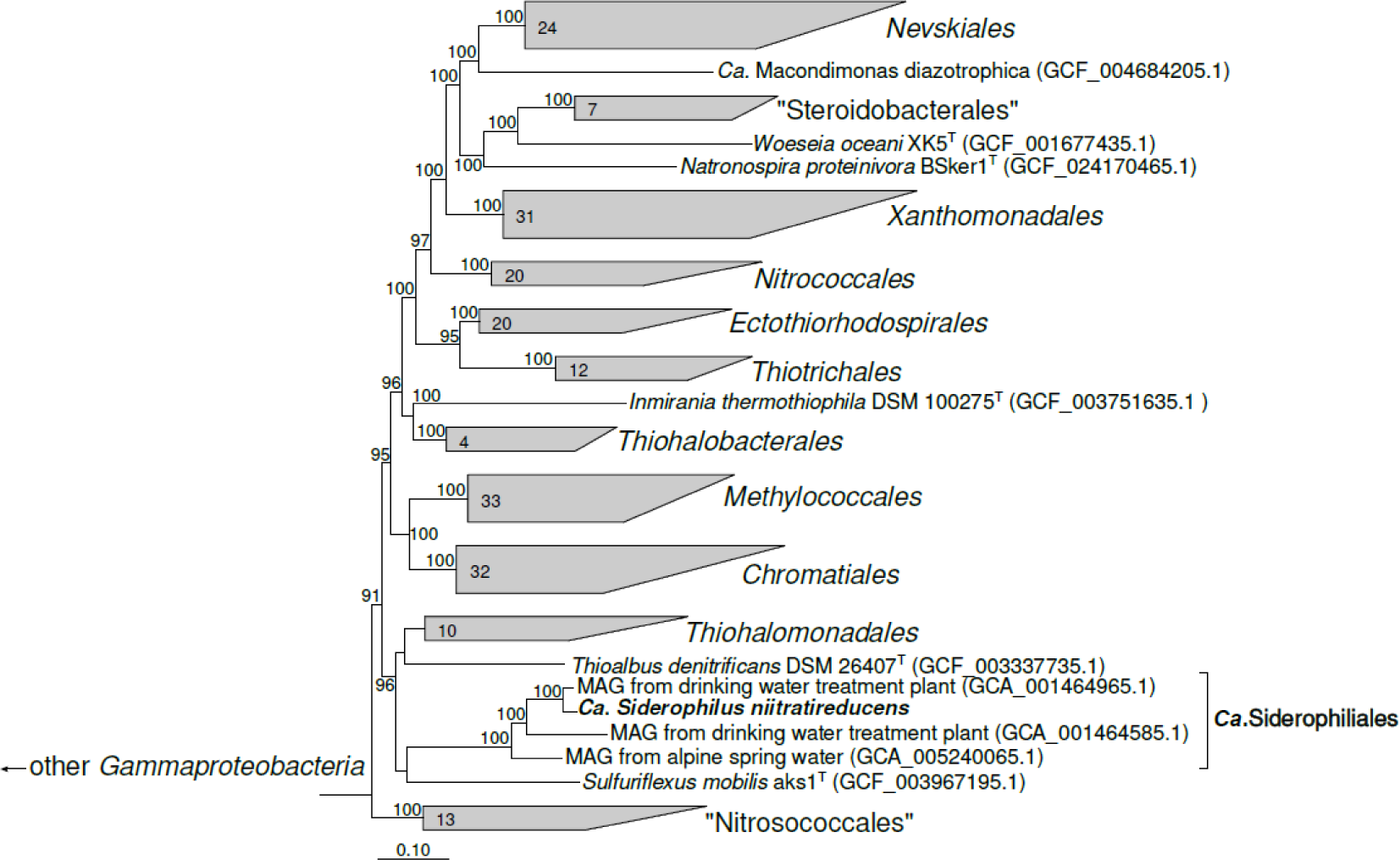
Phylogenetic position of the “Ca. Siderophilus niitratireducens” based on sequence analyses of concatenated alignment of 120 single-copy conserved bacterial protein markers (49) - taxonomic designations correspond to the Genome Taxonomy DataBase 207). The trees were built using IQ-TREE2 (ref. 23) with approximate likelihood-ratio test for branches (62). Bootstrap consensus tree is shown with values above 90% placed at the nodes. Bar, 0.10 changes per position.

### 2.4. *Autotrophy in “Ca.* Siderophilus nitratireducens*”*

To resolve the main anabolic and catabolic pathways of “*Ca*. Siderophilus nitratireducens”, open reading frames (ORF) were predicted and annotated (Table 3, detailed version in Table S1). The genome contains marker genes coding for two key proteins of autotrophic CO_2_ fixation via the reductive pentose phosphate (Calvin–Benson–Bassham; CBB) cycle, including the large and small subunits of the ribulose-1,5-bisphosphate carboxylase-oxygenase (*rbc*LS form I) and the phosphoribulokinase (*prk*). Genes encoding for carboxysomal shell proteins and carbonic anhydrase were also present, further supporting the inorganic carbon uptake ability of “*Ca.* Siderophilus nitratireducens*”*. The absence of phosphofructokinase (*pfk*) indicates a modified glycolytic pathway initiating at the glyceraldehyde 3-phosphate level. All tricarboxylic acid (TCA) cycle genes were identified except for fumarate hydratase (*fh*). However, the glyoxylate shunt enzymes malate synthase (*glc*B) and isocitrate lyase (*ace*A) were present. Taken together, these findings suggest the capability for full autotrophic growth of “*Ca.* Siderophilus nitratireducens*”*.

### 2.5. Iron oxidation in “Ca. Siderophilus nitratireducens”

The presence of a monoheme *c* cytochrome *cyc2*, a primary iron oxidation gene, suggests that “*Ca.* Siderophilus nitratireducens*”* can use Fe^2+^ as an electron donor. Other common Fe^2+^ oxidases, namely the diheme *c* cytochrome cyc1 and the multiheme c cytochromes MtoA and MtoB, were not annotated. Despite the close phylogenetic proximity to lithoautotrophic sulfur-oxidizing bacteria, the genes of sulfide dehydrogenases Sqr and FccAB and sulfite dehydrogenases SorAB and SoeABC were not identified.

In terms of potential catabolic electron acceptors, the genes for a periplasmic nitrate reductase (*nap*ABCD and its membrane ferrodoxins *nap*GH) and a *cbb*_3_-type cytochrome c oxidase (*cco*NOP) were annotated. However, genes encoding for other known denitrification reductases, namely membrane-bound nitrate reductase (*nar*GHI) and nitrite, nitric oxide and nitrous oxide reductases, *nir*K/*nir*S, *nor*BC and *nos*Z respectively, were not found (Table 3, detailed version in Table S2). Additionally, alternative oxidases, such as the cytochrome *bd* ubiquinol oxidase (*cyd*AB) or the *aa_3_*-type cytochrome c oxidase (*cox*ABCD) were not identified. Also, genes of dissimilatory sulfate reduction (*apr*AB and *dsr*ABC) and the *sox* complex, responsible for sulfate reduction could not be identified. These findings suggest that “*Ca.* Siderophilus *nitratireducens”* relies exclusively on nitrate and oxygen as electron acceptors.

### 2.6. Complete denitrification: a collaborative effort of iron-oxidizing autotrophs and organoheterotrophs

Iron oxidation genes were identified in five MAGs, namely MAG.13 (“*Ca.* Siderophilus nitratireducens*”*), MAG.29 (g_*Rhizobacter*) and MAG.26, MAG.34 and MAG.16 (f_*Gallionellaceae,* commonly associated with autotrophic iron oxidation). These MAGs also encoded for the central enzymes of the carbon dioxide fixation via the CBB cycle. While all 13 MAGs contained genes encoding for at least one denitrification enzyme (Table 2), none of them possessed a comprehensive gene set to fully reduce nitrate to dinitrogen gas. Dissimilatory nitrate reduction to nitrite, the first denitrification step, was present in 9 MAGs, while the final step, nitrous oxide reduction to nitrogen gas, was only found in MAG.26 (g_*Rhizobacter*), MAG.18 (f_*Anaeromyxobacteraceae*) and MAG.10 (f_*Chitinophagaceae*). The five most abundant MAGs (> 5 %) alone accounted for up to 50 % of the community and covered the full denitrification process (Figure 3). Interestingly, two distinct potential niches were identified. The autotrophic iron oxidizers, *“Ca.* Siderophilus *nitratireducens*” and MAG.26 (f_*Gallionelleceae*), performed the initial denitrification reductions, while the lower-abundant organoheterotrophs complemented the reduction of (at least) NO to nitrous oxide and, possibly took advantage from the autotrophically fixed carbon excreted by the iron oxidizers. Noticeably, due to the absence of sufficient biodegradable organic matter in the influent, a portion of the biologically generated NO must have been chemically reduced to N_2_O with Fe^2+^(20).

**Figure 3.**
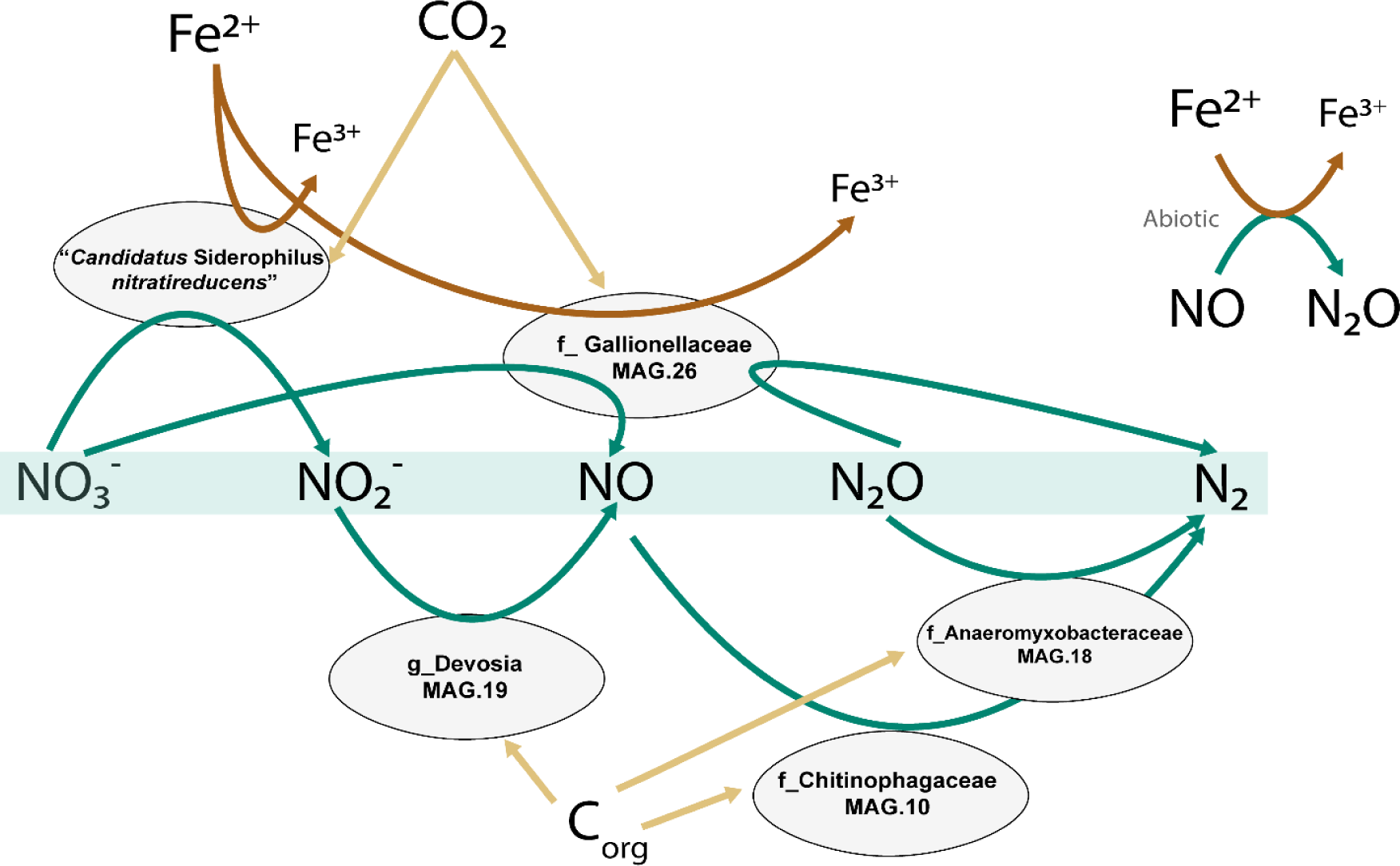
Genome-based conceptual model of substrates fluxes within the microbial community represented by the five most abundant MAGs. Putative autotrophic iron oxidizers perform the upstream part of denitrification, while flanking communities reduce the toxic intermediates to innocuous dinitrogen gas. The putative autotrophic metabolism was inferred based on the presence of ribulose-1,5-biphosphate carboxylase/oxygenase (RuBisCO) and phosphofructokinase (pfk).

**Table 2.**
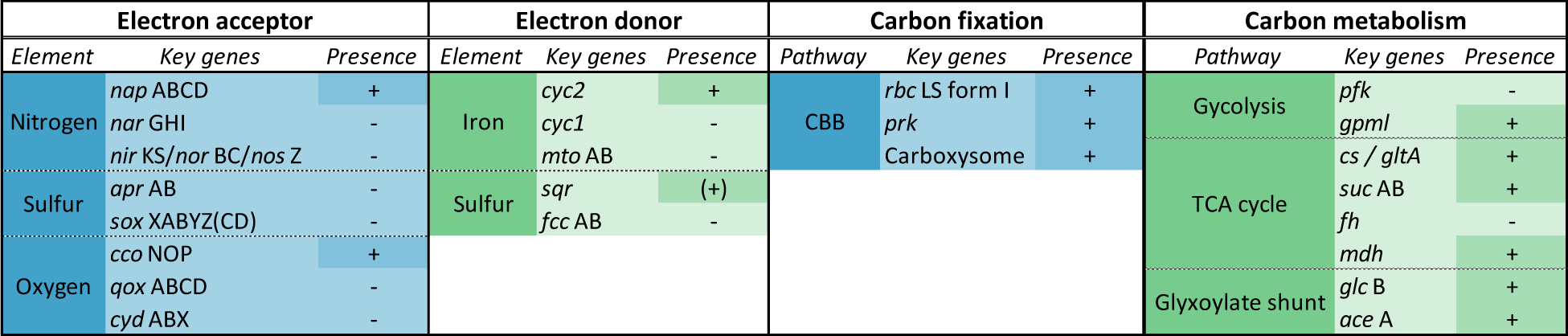
Key enzyme of the main catabolic and anabolic pathways of “Ca. Siderophilus nitratireducens”. “+” and “-“ indicate presence or absence in the genome. CBB, Calvin-Benson-Bassham cycle; TCA, Tricarboxylic acid cycle/Krebs cycle.

## 3. Discussion

We established a pilot-scale filter on nitrate-rich groundwater to elucidate the structure and metabolism of nitrate-reducing iron-oxidizing microbial communities under oligotrophic conditions mimicking natural groundwater. The enriched community stoichiometrically removed iron and nitrate, and was dominated by a genome belonging to a new *Candidate* order, named Siderophiliales. The genome of this new species, “*Ca.* Siderophilus *nitratireducens*”, encoded the genes for iron oxidation (cytochrome *c cyc2*) and the periplasmic nitrate reductase (*nap*), thereby supporting the hypothesis of nitrate-driven iron oxidation being its primary metabolism under the restricted availability of alternative substrates. The absence of other denitrification genes suggests a short catabolic path, which might offer a kinetic advantage in terms of higher iron oxidation rates (21) especially under N limitation (22). “*Ca.* Siderophilus *nitratireducens*” was identified as a putative-autotroph, adding the additional challenge of energy and electrons needs for anabolic CO_2_ fixation to the growth on iron, a weak electron-donor at standard conditions (14). However, thermodynamic evaluations do suggest, that *nap*-dependent iron oxidation can sustain growth, given realistic Fe^3+^/Fe^2+^ and 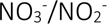 concentration ratios (SI 4). An essential factor in this context is the quasi-instantaneous precipitation of the biologically formed Fe^3+^ as iron oxides under circum-neutral pH (23). Biochemically, by accepting electrons from the quinol pool, *nap* consumes cytoplasmic protons effectively generating a proton gradient. While the specific mechanism by which this thermodynamic potential is harnessed for carbon fixation remains to be fully elucidated, to the best of our knowledge this is the first report of *nap*-based lithoautotrophic growth. The presence of the sole *nap* also broadens the biochemical diversity of NRFOs currently believed to rely exclusively on the membrane-bound nitrate reductase (*nar*) (9,24–27).

The subsequent reduction of the produced nitrite resulted from the concerted activity of putative autotrophic iron-oxidizers and organoheterotrophs. Within the microbial community, the second most abundant genome, MAG.26 (f_*Gallionellaceaea*), featured the genetic potential for iron oxidation and most denitrification steps, with the exception of nitrous oxide reductase (*nor*). Interestingly, this genome contained genes for CO_2_ fixation, a trait mirrored in all other less abundant genomes with the ability to oxidize iron. This suggests that autotrophy may represent an essential trait for NRFOs in anoxic groundwaters where the dissolved organic carbon is largely non-biodegradable (17). The three second most abundant genomes, MAG.18 (f_*Anaeromyxobacteraceae*), MAG.19 (g_*Devosia*) and MAG.10 (f_*Chitinophagaceae*) were found to lack the genes for iron oxidation and CO_2_ assimilation. Yet, these genomes encompassed the full denitrification pathway starting from nitrite. Besides the likely occurrence of chemical reduction of NO to N_2_O (28), we speculate that these heterotrophs complemented the NRFOs for at least the reduction of NO using autotrophically fixed organic carbon as substrate. A similar metabolic network was also recently observed in mesophilic NRFO communities (12). Overall, the measured iron and nitrate consumption yield of 7.1 mol Fe^2+^: mol 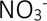 is consistent with the expected 5.6, *i.e.* considering the theoretical catabolism (eq.1) and the recently estimated 12 % of electrons used for growth (29), but higher than the experimentally observed range of 3.8 - 4.7 (7,24,30). At first, we hypothesized nitrate ammonification to be the reason for the slight excess in iron oxidation, yet none of the putative iron-oxidizing genomes encoded for the common *nrf* nor for the newly reported octaheme complex (31). Also, the oxygen sporadically detected in the influent was always significantly below the quantification limit of 3 μM, a conservative concentration that alone would explain less than 15 % of the total iron consumption via chemical oxidation. As no Fe^3+^ was detected in the reactor effluent, all iron necessarily accumulated inside the reactor either as Fe^2+^ or Fe^3+^ precipitates. X-ray diffraction and Mossbauer spectroscopy identified over 94% of the Fe in solids as amorphous ferrihydrite, an Fe^3+^ oxide, with less than 6% of the solids attributed to magnetite, an Fe^2+^-Fe^3+^ oxide typically formed under anaerobic conditions (SI 5 and 6). Consequently, the Fe^2+^ unaccounted for was likely continuously adsorbed onto the newly-formed Fe^3+^ oxides, a well-studied phenomenon (32), yet the extent to which this occurred was not investigated. In conclusion, pending experimental validation, we surmise that NRFO microorganisms may not only contribute to iron removal by direct oxidation but also by continuously providing newly-formed iron oxides for its adsorption.

*Description of “Ca.* Siderophilus nitratireducens” gen. nov., sp. nov.

*Siderophilus* (Si.de.ro’phi.lus Gr. masc.n. *sidêros* iron; Gr. masc. adj. *philos* loving; N.L. masc.

n. *Siderophilus* (loving iron).

ni.tra.ti.re.du’.cens (N.L. masc. n. *nitras (gen. nitratis)*, nitrate; L. pres. part. *reducens*, converting to a different state; N.L. part. adj. *nitratireducens*, reducing nitrate).

Autotrophic nitrate-reducing iron-oxidizing bacterium isolated from a filtration unit fed with anaerobic groundwater with iron(II) and nitrate.

## 4. Materials and Methods

### 4.1. Groundwater and pilot-scale filter characteristics

An iron reducing microbial community was enriched anoxically on the granular activated carbon of a 10-L pilot-scale filter in Emmen (the Netherlands) (Figure S1, Figure S2 and Table S1). The media was devoid of any previously formed biofilm. The anoxic, nitrate-rich groundwater (−75,2 ± 28.4 mV) featured constant Fe^2+^ and 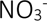 concentrations, 236 ± 4 µM and 8.1 ± 2.1 µM respectively (Table 3). Oxygen was consistently below quantification limit (3 μM). The groundwater pH and temperature were 6.7 ± 0.2 and 10.5 ± 0.1 °C, respectively. The filter was operated at a filtration flowrate of 3.8 m·h^−1^ during 120 days. After 75 days of steady-state operation, the influent nitrate concentration was manually increased in four steps up to 83.8 ± 0.6 µM, when the system changed from nitrate (NO_3_^−^ < 1 µM) to iron limiting conditions (Fe^2+^ < 4 µM).

**Table 3.** Influent and effluent water characteristics corresponding to average and standard deviation of daily measurements of days 21-77, during the nitrate-limited steady-state. Fe^3+^ was calculated as described in the following section.

| Parameter | Units | Influent Value | Effluent Value |
| --- | --- | --- | --- |
| pH | - | $6.9 \pm 0.4$ | $6.7 \pm 0.1$ |
| T | $^{\circ}C$ | $10.5 \pm 0.1$ | $10.9 \pm 0.6$ |
| ORP | mV | $-64 \pm 17$ | $-58 \pm 18$ |
| $O_2$ | $\mu mol \cdot L^{-1}$ | $<3^*$ | $<3^*$ |
| $NH_4^+$ | $\mu mol \cdot L^{-1}$ | $11 \pm 1.0$ | $11 \pm 8.1$ |
| $NO_2^-$ | $\mu mol \cdot L^{-1}$ | $<0.2^*$ | $<0.2^*$ |
| $NO_3^-$ | $\mu mol \cdot L^{-1}$ | $8.1 \pm 2.1$ | $<1$ |
| $Fe^{2+}$ | $\mu mol \cdot L^{-1}$ | $236 \pm 4$ | $178 \pm 5$ |
| $Fe^{3+}$ | $\mu mol \cdot L^{-1}$ | $2 \cdot 10^{-12}$ | - |
| DOC | $mg \cdot L^{-1}$ | $3.1 \pm 0.1$ | $2.0 \pm 0.2$ |

\* Below detection limit

### 4.2. Dissolved Fe^3+^ estimation

At pH 7, Fe^3+^ has a markedly low solubility and precipitates as iron oxyhydroxide (Fe(OH)_3_). Thermodynamically, this phase transition favours the oxidation of Fe^2+^ to Fe^3+^, and the resulting low Fe^3+^ concentration is the primary driving force of eq. 1 (ref. 36). In our filter, the dissolved concentration of the Fe^3+^, resulting from Fe^2+^ oxidation, was always below detection limit (0.01mg·l^−1^). To discuss the thermodynamic feasibility of the NRFO process, we estimated the steady-state [Fe^3+^]/[Fe^2+^] ratio following the method proposed by Gorski et al. (2016), which assumes thermodynamic equilibrium between [Fe^2+^] – [Fe^3+^] – [FeOx] phases based on the fact that the hydroxylation of dissolved Fe^3+^ is quasi-instantaneous at pH > 3 (ref. 37). Consequently, the following equation can be used to determine the Fe^3+^ concentration as function of pH and the solid solubility constant.

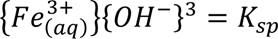

The most abundant iron oxide in the sand filter was amorphous ferrihydrite (SI 5 and 6), with a K_sp_ of 10^−39^ (35). Therefore:

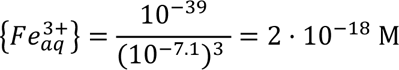

### 4.3. Analytic procedures

Samples for ammonium, nitrite, and nitrate quantification were immediately filtered through a 0.2 µm nanopore filter and measured within 12 h using photometric analysis (Gallery Discrete Analyzer, Thermo Fischer Scientific, Waltham, Massachusetts, USA). Samples for dissolved iron were filtered through a 0.2 μm nanopore filter and quantified by ICP-MS (Analytik Jena, Jena, Germany). Temperature, pH, oxidation-reduction potential (ORP), and dissolved oxygen concentration (DO) were monitored daily using a HI9829-01042 multiparameter analyser (Hanna Instruments, Smithfield, RI, USA) in the raw water, after nitrate dosage and in the effluent.

### 4.4. DNA extraction and quality control

Nucleic acid extraction was carried out using DNeasy PowerSoil Pro Kit (QIAGEN, Hilden, Germnay) following manufacturer instructions. To improve DNA recovery and avoid the interference of carbon with the extraction, 25 µL of 20 g·l^−1^ autoclaved (20 min, 121° Celsius, 2 bar) skimmed milk solution (Sigma Aldrich, Saint Louis, MO, USA) were added to the extraction tube. After extraction, DNA was concentrated to 7.68 ngDNA·µ^−1^ using Microcon centrifugal filter units YM-100 (MilliporeSigma, Burlington, MA, USA) following the manufacturer’s instructions. DNA was quantified with the Qubit 4 Fluorometer and Qubit dsDNA HS assay kit (Invitrogen, Waltham, MA, USA) following the manufacturer’s instructions. DNA purity was determined using a NanoDrop One Spectrophotometer (Thermo Fisher Scientific, Waltham, MA, USA).

### 4.5. Library preparation, sequencing and reads processing

Long-read and short-read DNA sequencing were carried out independently. Long-read library preparation was carried out using the ligation sequencing kit SQK-LSK 109 (Oxford Nanopore Technologies, Oxford, UK). R.9.4.1 flowcells on a GridION were used for sequencing. Raw data was basecalled in super-accurate mode using Guppy v.5.0.16 (https://nanoporetech.com). Raw reads were quality-filtered and trimmed using Filtlong (https://github.com/rrwick/Filtlong) to remove reads below 4000 kb and mean quality score below 80. Adapters were removed using Porechop v.0.2.3 (https://github.com/rrwick/Porechop).

Short-read library preparation was performed using the Nextera XT kit (Illumina, San Diego, CA, U.S.A.) according to the manufacturer’s instructions. The libraries were pooled, denatured, and sequenced with Illumina MiSeq (San Diego, CA, USA). Paired end sequencing of 2 x 300 base pairs was performed using the MiSeq Reagent Kit v3 (San Diego, CA, USA) according to the manufacturer’s instructions. Raw sequencing data was quality-filtered and trimmed using Trimmomatic v0.39 (HEADCROP:16 LEADING:3 TRAILING:5 SLIDINGWINDOW:4:10 CROP:240 MINLEN:35) (36). Sequencing data quality was analyzed using FastQC v0.11.7 before and after trimming (37).

### 4.6. Reads assembly and binning

Reads assembly and binning were done as in (38) with minor modifications. Long-reads were assembled using Flye v. 2.9-b1768 (39) with the ‘–meta’ setting enabled and the ‘–nano-hq’ option. Polishing was carried out with Minimap2 v.2.17 (40), Racon v. 1.3.3 (41), Medaka v.1.4.4 (two rounds) (https://github.com/nanoporetech/medaka). At the end, short-reads were incorporated in a final round of polishing with Racon. Both long- and short-raw reads were independently mapped back to the assembled contigs using BWA-MEM2 (42). SAMtools v1.14 was used to determine contig coverage and for indexing with default settings (43).

Automated binning was carried out with the long-reads assembly (polished with short-reads) using MetaBAT2 v. 2.12.1 (ref. 46) with ‘-s 500000’, MaxBin2 v. 2.2.7 (ref. 47), and Vamb v. 3.0.2 (ref. 48) with ‘-o C–minfasta 500000’. Additionally, contig coverage from the short-reads assembly was provided as input to the three binners to improve binning. Output integration and refinement was done in DAS Tool v. 1.1.2 (ref. 49). CoverM v. 0.6.1 (https://github.com/wwood/CoverM) was applied to calculate the bin coverage (using the ‘-m mean’ setting) and the relative abundance (‘-m relative_abundance’). Additional manual bin polishing was done in R using mmgenome (https://github.com/MadsAlbertsen/mmgenome).

### 4.7. Assembly processing and gene annotation

The completeness and contamination of the genome bins were estimated using CheckM v. 1.1.2 (ref. 50). The bins were classified using GDTB-Tk v. 1.5.0 (ref. 22) 202 database. Barrnap v 0.9 (https://github.com/tseemann/barrnap) and structRNAfinder (50) were used to predict 23S, 16S and 5S ribosomal RNA sequences, and transfer RNA sequences were determined using tRNAscan-SE v.20 (51) with *default* search mode. Bins were classified using the Minimum Information about a Metagenome-Assembled Genome (MIMAG) standards (18): high-quality bins were > 90 % complete and < 5 % contaminated, and contained full-length 23S, 16S and 5S ribosomal RNA genes and ≥ 18 transfer RNA genes. Bins with completeness > 50 % and contamination < 10 % were classified as medium-quality, and bins with completeness <50% and <10% contamination as low-quality bins. The remaining ones were considered contamination. 68% of the filtered reads rendered high- and medium-quality bins, 11% low-quality bins and 21.1% were unbinned.

The open reading frames (ORF) of the 10 resulting high-quality and 3 medium-quality bins with relative abundance > 0.5 % were predicted using Prodigal v2.6.3 (ref. 53) and functionally annotated with GhostKoala v2.2 (53) (Kyota Enciclopedia of Genes and Genomes; accessed March 2022). FeGenie (54) was used to improve the annotation of the iron metabolism using the metagenomics (‘-meta’) settings. To refine the annotation for MAG.13 (*Candidatus* Siderophilus nitratireducens), the genome was uploaded to the National Center for Biotechnology Information (NCBI) database Prokaryotic Genome Annotation Pipeline v6.1 (ref. 54). Additionally, manual annotation of genes potentially relevant but not automatically annotated was done by aligning a set of manually selected sequences from UniProtKB against the translated ORFs from MAG.13 with local blastp v2.13 (ref. 55). After annotation, all the predicted genes of interest (manually and automatically annotated) were translated and aligned against the Non-redundant protein sequences (nr) database from NCBI using blastp (accessed June 2022) and accepted only if the coverage was > 70 % (54) and the identity > 35 % (57).

RStudio v1.4.1106 was used for data analysis and visualization.

### 4.8. Phylogenetic tree construction

Genome-based phylogenetic reconstruction was done by using 120 bacterial single copy conservative marker genes, as described previously (58). The trees were built using the IQ-TREE 2 (ref. 23) with fast model selection via ModelFinder (60) and ultrafast approximation for phylogenetic bootstrap (61), as well as approximate likelihood-ratio test for branches (62). Whole genome comparison was conducted by using two different methods: Average Nucleotide Identity (ANI), using JSpeciesWS web server and DNA-DNA Hybridization (DDH) by the Genome-to-Genome Distance Calculator 2.1 online tool (https://ggdc.dsmz.de/ggdc.php) (63).

## 5. Competing interest

The authors declare no competing financial interests

## 6. Acknowledgements

The authors would like to thank Dimitry Sorokin (TU Delft and Russian Academy of Sciences) for the thorough discussions and invaluable insights, Martin Pabst and Dita Heikens (TU Delft) for the proteomic data, and Mantas Sereika, Thomas Nielsen and Mads Albertsen (Aalborg University) for their help with Nanopore Sequencing and genome assembly and binning.

This work was financed by the NWO partnership program ‘Dunea–Vitens: Sand Filtration’ (project 17830) of the Dutch Research Council (NWO) and the drinking water companies Vitens NV and Dunea Duin & Water. ML was supported by NWO (VI.Veni.192.252).

## 7. Data availability statement

Raw reads and MAGs have been deposited in the National Center for Biotechnology Information (NCBI) website (https://www.ncbi.nlm.nih.gov/bioproject/) under BioProject PRJNA834785. The BioSample accession numbers for the raw reads and the five most abundant MAGs are SAMN28600298, SAMN28058410 (“*Ca*. Siderophilus nitratireducens), SAMN36381704 (*f_Gallionelaceae*), SAMN36381891 (*f_Anaeromyxobacteraceae*), SAMN36382736 (*g_Devosia*), SAMN36401011 (*f_Chitinophagaceae*).

## 8. Authors contributions

FCR, JB, MvL, DvH and ML conceived the study. JB and SD designed the filter and performed the on-site experiments. FCR run the metagenomics analysis, and AM the phylogenetic classification. FCR and GS conducted the thermodynamics calculations. SM performed the solid characterization. FCR, GS, MvL, DvH and ML performed data analysis and/or helped interpret the results. FCR wrote the manuscript, with contributions from GS, MvL, DvH and ML. All co-authors critically reviewed the manuscript and approved the final version.

## Notes

### Competing Interest Statement

The authors have declared no competing interest.

