## Supplementary Material for "“*Candidatus* Siderophilus nitratireducens”: a psychrophilic, *nap*-dependent nitrate-reducing iron oxidizer within the new order Siderophiliales"

### 1. Pilot-scale filter and groundwater characteristics

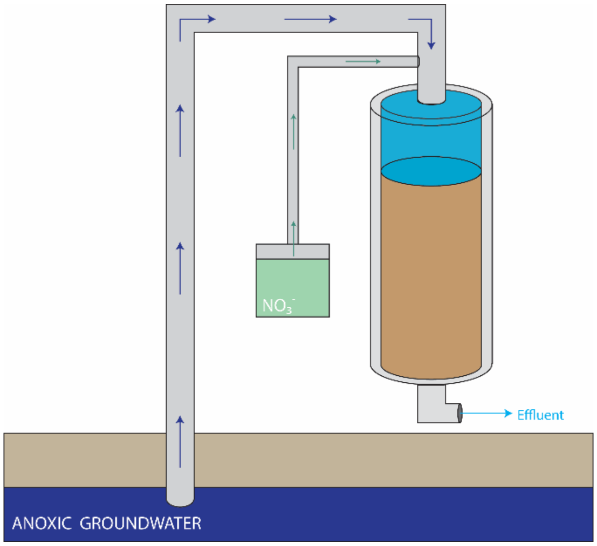


*Figure S1. Scheme of the pilot-scale set-up. Anoxic groundwater containing iron and nitrate was fed into a filter filled with granular activated carbon. Nitrate was added manually into the groundwater prior to the filter in the second phase of the experiment.*

*Table S1. Operational and design parameters of the pilot-scale filter.*

| Parameter | Units | Value |
|---|---|---|
| Bed height | m | 1.34 |
| Filter area | $m^2$ | 0.0078 |
| Empty bed contact | min | 21 |
| Filtration velocity | m/h | 3.8 |

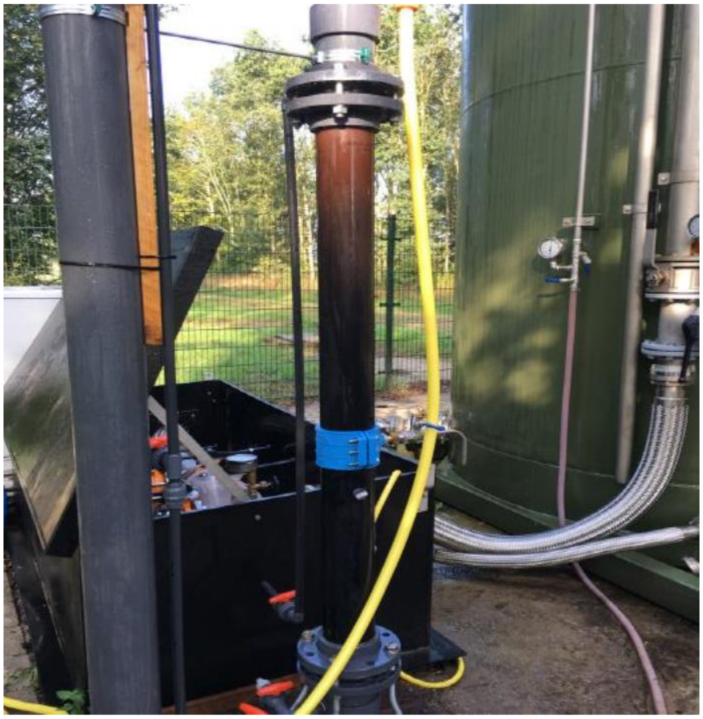

*Figure S2. Pilot-scale rapid sand filter filled with granular activated carbon and fed with anoxic groundwater.*

#### 2. Continuous operation of pilot-scale filter

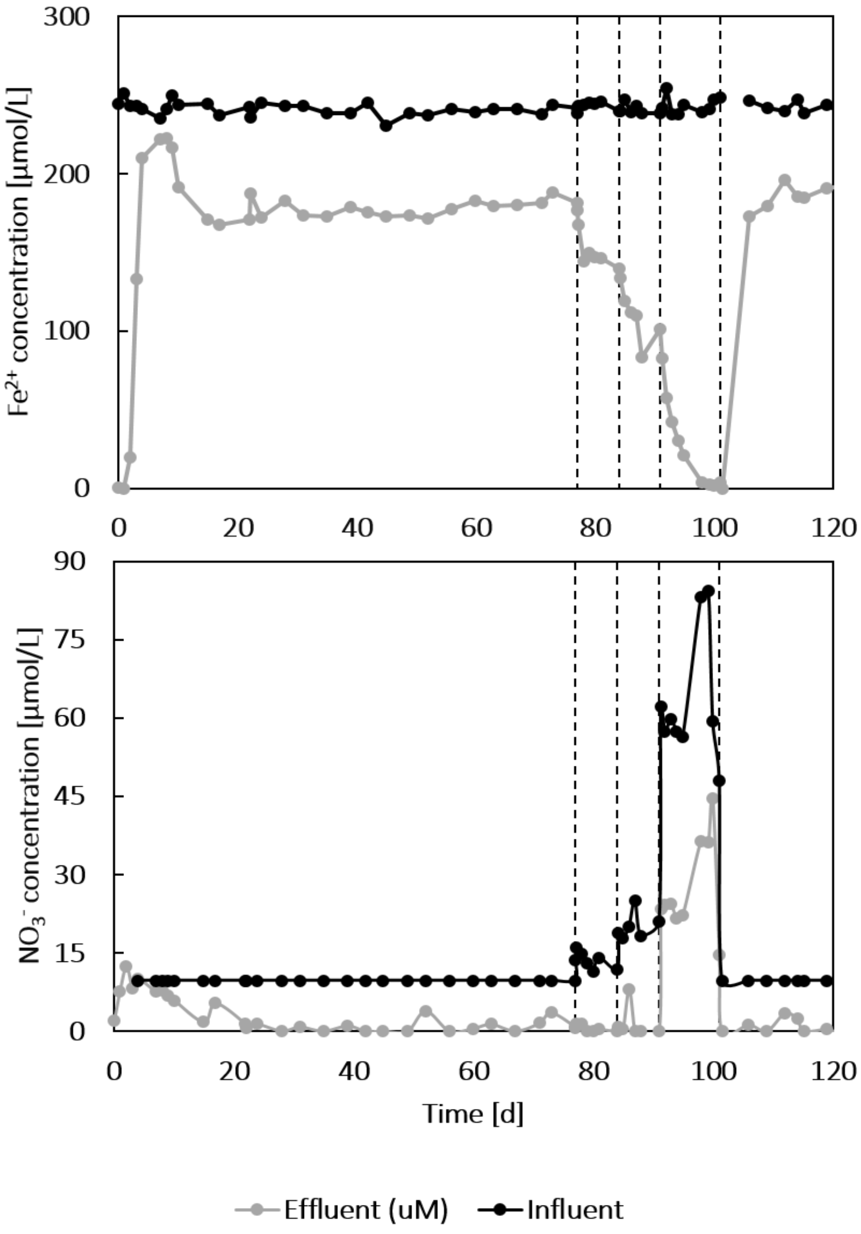


*Figure S3. Influent and effluent nitrate (top) and $Fe^{2+}$ (bottom) concentration in the groundwater-fed pilot-scale sand filter along 120 days of continuous operation from. The concentration of $Fe^{2+}$ in groundwater was constant throughout the experiment (236 ± 4 mM). Nitrate was dosed in the influent to step-wise increase the natural groundwater concentration (8.1 ± 2.1 up to 83.8 ± 0.6 mM) and confirm the nitrate-reducing iron-oxidizing activity. Nitrate is completely consumed except in the period between days 91 and 101, during which $Fe^{2+}$ was limiting.*

#### 3. Metagenomic analysis

*Table S2. Main metabolic pathways and enzymes in the genome of "Ca.* Siderophilus *Nitratireducens". ORF were predicted using Prodigal v2.6.3.* (1)*, and functional annotation was carried out with GhostKoala v.2.2 (ac. March 2022)* (2) *and the Prokaryotic Genome Annotation Pipeline v6.1.* (3)*. The annotation of iron oxidation genes was refined with FeGenie* (4)*. Predicted genes were aligned against the Non-redundant proteins sequences (nr) database from NCBI using blastp and accepted if coverage >70% and identity > 35%* (5)*.*

*[See Table as attached separate Excel file]*

#### 4. On the thermodynamic feasibility of Nap-dependent nitrate-reducing iron-oxidation

Based on its genome, "*Ca.* Siderophilus *nitratireducens*" is potentially an autotrophic organism that obtains energy by oxidizing $Fe^{2+}$ to $Fe^{3+}$ while reducing $NO_3^-$ to $NO_2^-$.Due to iron being a weak electron donor, the standard Gibbs free energy of the reaction is positive under standard biological conditions (pH 7) and thus the process is therefore thermodynamically unfavourable:

$$Fe^{2+} + 0.5NO_3^- + H^+ \rightarrow Fe^{3+} + 0.5NO_2^- + 0.5H_2O \quad \Delta G^{01} = 35.2\frac{kJ}{e\text{-}mol}$$

The exceptionally low solubility constants of iron oxides, ranging from $10^{-34}$ to $10^{-42}$ (6) result in very low $Fe^{3+}$ concentrations, thereby precipitation provides a thermodynamic driving force for the oxidation of $Fe^{2+}$ to $Fe^{3+}$ (7). Under our specific operational conditions (283 K, < 1 µmol$NO_3^- \cdot l^{-1}$, < 0.2 µmol$NO_2^- \cdot l^{-1}$), the reaction becomes favourable owing to the calculated $Fe^{3+}/Fe^{2+}$ ratio in the order of $10^{-16}$ (conservative value; see SI 2):

$$\Delta G^1 = \Delta G_r^{01} + RTln\left[\frac{[Fe^{3+}][NO_2^-]^{\frac{1}{2}}}{[Fe^{2+}][NO_3^-]^{\frac{1}{2}}}\right] = -66.2\frac{kJ}{e\text{-}mol}$$

A Gibbs free energy of -66 kJ/e-mol is in principle enough to net transport three to four protons over the cytoplasmic membrane, *i.e.* 15 kJ·mol$^{-1}$ (8), and would yield at least one ATP. If other iron precipitation products are formed, the redox potential of $Fe^{2+}/Fe^{3+}$ increases further. Even at more conservative $Fe^{3+}/Fe^{2+}$ ratios and pH, due to non-equilibrium conditions and product gradients, the biological process remains favourable (see Figure S4).

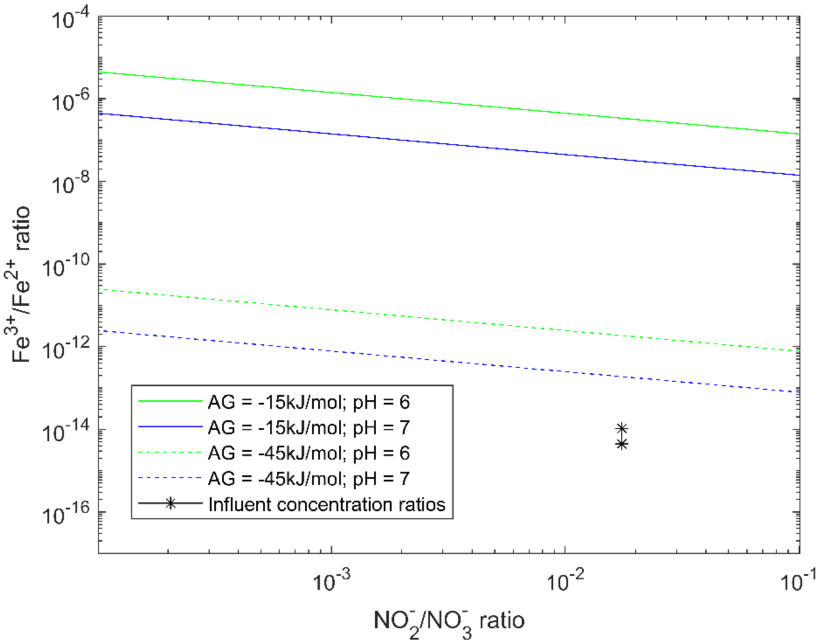


*Figure S4. Gibbs energy dissipated during nitrate-reducing iron oxidation as function of the ratio of substrates and products. As reference, lines represent the ratios at which the minimum energy required to generate* pmf *(-15 kJ·mol$^{-1}$, full lines) and ATP (-45 kJ·mol$^{-1}$, dashed lines) is generated at intracellular pH of 6 (green) or 7 (blue) (Müller & Hess, 2017). Star dots represent the actual ratios based on measured ($NO_2^-/NO_3^-/Fe^{2+}$) and calculated ($Fe^{3+}$) influent reactor concentrations. Note that $NO_2^-$ was below detection limit (< 0.2 µmol·l$^{-1}$), so that the actual reactor conditions were likely even more favorable, i.e. more on the left .*

### 5. Solid identification with Mössbauer spectroscopy

#### *5.1. Materials and methods*

Transmission $^{57}$Fe Mössbauer spectrum was collected at 4.2 K with a sinusoidal velocity spectrometer using a $^{57}$Co(Rh) source. Velocity calibration was carried out using an α-Fe foil at room temperature. The source and the absorbing sample were kept at the same temperature during the measurement. The Mössbauer spectrum was fitted using the Mosswinn 4.0 program (9).

#### *5.2. Results*

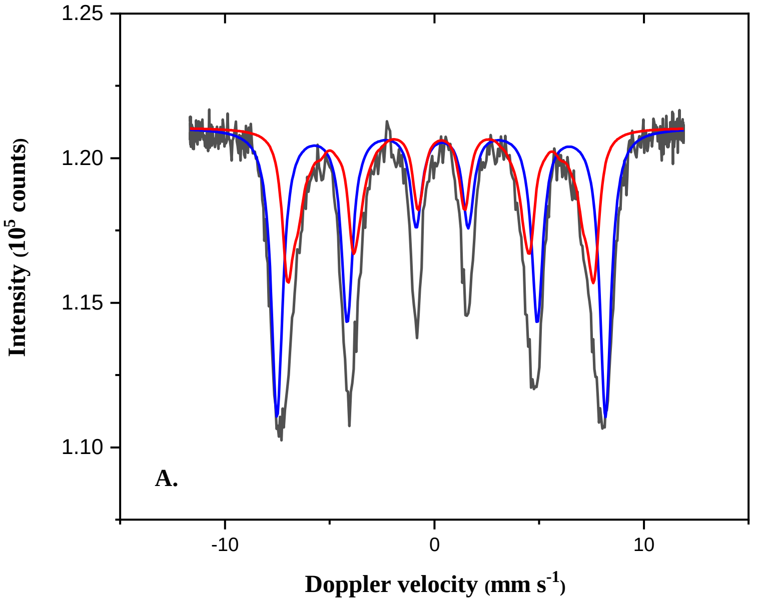


*Figure S5. Mossbauer spectra obtained at 4.2 K with iron oxide minerals.*

*Table S3.* The Mössbauer fitted parameters of iron oxide minerals

| *Sample* | *IS* (mm·s$^{-1}$) | *QS* (mm·s$^{-1}$) | *Hyperfine field (T)* | *Γ* (mm·s$^{-1}$) | *Phase* | *Spectral contribution (%)* |
|---|---|---|---|---|---|---|
| B. NDFO | 0.32 | -0.04 | 43.1* | 0.56 | $Fe^{3+}$ (Ferrihydrite[a]) | 43 |
| | 0.35 | -0.05 | 48.7 | 0.65 | $Fe^{3+}$ (Ferrihydrite[b]) | 57 |

*Experimental uncertainties: Isomer shift: I.S. ± 0.01 mm s-1; Quadrupole splitting: Q.S. ± 0.01 mm s-1; Line width: Γ ± 0.01 mm s-1; Spectral contribution: ± 3%; *Average magnetic field; a,bFerrihydrite structures with different crystallinity degrees.*

#### 6. Solid identification with X-ray diffraction

*[See enclosed document with XRD spectra]*
